# A novel feature selection for RNA-seq analysis

**DOI:** 10.1101/209841

**Authors:** Henry Han

**Affiliations:** Department of Computer and Information Science Fordham University, Lincon Center, New York, NY 10023

**Author notes:** Email address (Henry Han).

**Keywords:** RNA-seq, Feature selection, differential expression analysis, 2010 MSC: 00-01, 99-00

## Abstract

RNA-seq data are challenging existing omics data analytics for its volume and complexity. Although quite a few computational models were proposed from different standing points to conduct differential expression (D.E.) analysis, almost all these methods do not provide a rigorous feature selection for high-dimensional RNA-seq count data. Instead, most or even all genes are invited into differential calls no matter they have real contributions to data variations or not. Thus, it would inevitably affect the robustness of D.E. analysis and lead to the increase of false positive ratios.

In this study, we presented a novel feature selection method: nonnegative singular value approximation (NSVA) to enhance RNA-seq differential expression analysis by taking advantage of RNA-seq count data’s non-negativity. As a variance-based feature selection method, it selects genes according to its contribution to the first singular value direction of input data in a data-driven approach. It demonstrates robustness to depth bias and gene length bias in feature selection in comparison with its five peer methods. Combining with state-of-the-art RNA-seq differential expression analysis, it contributes to enhancing differential expression analysis by lowering false discovery rates caused by the biases. Furthermore, we demonstrated the effectiveness of the proposed feature selection by proposing a data-driven differential expression analysis: NSVA-seq, besides conducting network marker discovery.

## 1. Introduction

RNA-seq provides a revolutionary way to unveil transcription by using ultra-high-throughput sequencing technologies to generate hundreds of million short reads from RNA molecules [1, 2, 3]. As raw RNA-seq data, the short reads usually ask several to even hundreds of Gigabytes storage. The short reads are further assembled or aligned against a reference genome (e.g. human genome) to produce a transcriptome by using assembly or alignment tools such as Bowtie, SOAPdenovo-Trans, SOAP3, or HTSeq [4, 5, 6]. As a genome level transcription map, the transcriptome consist of the expression levels of all genes in transcription and each gene’s expression is represented as the number of short reads mapped to the gene in the alignment or assembly [7]. In fact, the terminology gene refers to more general biological features in transcription such as a gene, exon, or transcript [8, 9].

The transcription map can be represented by a nonnegative integer read count matrix 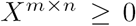 by collecting all read counts mapped to each gene, where each row and column represent a gene and sample respectively. According to different sources, a sample can be classified as a biological or technical replicate. The former is an alternative sequencing of a same biological sample, and the latter is the direct sequencing of an independent biological sample. For the convenience of description, we also use RNA-seq data to refer to the read count matrix *X* of the original RNA-seq data in this study.

RNA-seq data are actually biased data subject to sequencing depth and gene length by the nature of RNA-seq sequencing even after normalization [10, 11]. The sequencing depth bias means that those genes from a sample with a high sequencing depth, usually have higher expression levels (high counts) than the same genes from a sample with normal or low sequencing depth; The gene length bias refers to that long genes have more counts in RNA-sequencing than short genes. The biases bring challenges in normalization, differential expression (D.E.) analysis, and feature selection [10, 11].

However, they share the same characteristic with traditional omics data (e.g. microarray). That is, they both are high-dimensional data, i.e. the number of variables (genes) is generally much greater than the number of samples (observations) in a RNA-seq dataset 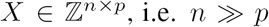. Unlike the traditional omics data, RNA-seq data are generally modeled by a poisson or negative binomial (NB) distribution instead of a normal or log-normal distribution [7, 8, 10].

An essential issue in RNA-seq analysis is to answer the query: ‘given a read count matrix, how to robustly determine whether the observed difference for a gene across two or more conditions is statistically significant?’. Quite a few differential expression (D.E.) analysis methods were proposed to answer it from different perspectives [8, 9, 11, 12, 13, 14, 15].

They can be categorized as either parametric or non-parametric methods according to whether they rely on statistical parameter estimation modeling approaches or not in D.E. analysis. The former assumes that RNA-seq data are subject to a well defined distribution and estimates corresponding parameters for the distribution before conducting hypothesis tests. For example, *DESeq* and *edgeR* methods both model RNA-seq data by the negative binomial (NB) distribution and estimate genes’ mean *µ* and variance *σ*^2^ parameters [8, 13]; The latter does not assume data are subject to any distribution. Instead, its differential expression call can be based on an empirical distribution of some statistics derived from the input data. For example, NOISeq finds differentially expressed (DE) genes by relying on two derived statistics: absolute-expression and log-fold change between conditions [11].

However, all the D.E. analysis methods usually invite almost all genes into D.E. analysis without conducting a rigorous feature selection for high-dimensional RNA-seq data. Although they usually employ some simple filtering techniques to exclude those genes with zero or low counts before normalization, such a low-count filtering is by no means a desirable feature selection for high-dimensional data. Especially, such a count-based filtering may have a risk to affect the following D.E. analysis by feeding it with many high-count genes due to sequencing depth and gene length biases. As a result, it not only would increase false positives in the D.E. analysis, but also easily lead to a misleading result that those genes are differentially expressed simply because they have higher coverages or long gene length.

As typical high-dimensional data, RNA-seq data call for a rigorous feature selection algorithm, which should be robust to the depth and gene-length biases, to select potentially D.E. genes for the sake of D.E. analysis, rather than its number of read counts only. Such a feature selection should overcome the weakness of the naive count-based filtering by removing redundant and noise-contained genes, instead of only low-count genes.

In fact, those high-count genes with very low variance values can be viewed as the redundant genes because they may not have real contributions to data variations. Furthermore, some low count genes with several exceptional high count peaks on few replicates in a same condition can be viewed as noise-contained genes, because the peaks could be generated from depth-based oversampling. The genes will be falsely identified as D.E. genes in differential calls, even if the observed differences between conditions are actually due to the artifacts of over-sequencing or library preparations instead of real reactions to a treatment.

On the other hand, a rigorous feature selection will strengthen RNA-seq normalization efforts to alleviate the effects of the depth and gene length bias factors by removing those genes not totally ‘corrected’ in normalization, in addition to lowering the computing complexities in the following D.E. analysis. As such, there needs a rigorous feature selection to glean meaningful genes to achieve a more targeted and accurate differential expression analysis.

In this study, we present a novel feature selection method: nonnegative singular value approximation (NSVA) to enhance RNA-seq analysis by taking advantages of RNA-seq data’s built-in non-negativity. The non-negativity is an important characteristic of RNA-seq data, but it is ignored in most feature selection methods. Nonnegatvity based analysis can contribute to enhancing data locality and capturing latent data behavior [20]. As a data driven feature selection, NSVA does not assume any priori distribution for RNA-seq data; As a variance-based feature selection, it selects genes according to its contribution to the first singular value direction of input data.

We compare the proposed feature selection method with its five peers in state-of-the-art RNA-seq differential expression analysis. It demonstrates robustness to depth bias and gene length bias in feature selection and contributes to more efficient D.E. analysis. To further explore NSVA’s effectiveness, we propose a data-driven differential expression analysis: NSVA-seq that is a novel nonparametric D.E. analysis without M-D odd ratio comparisons[11]. More importantly, it overcomes the weakness of existing D.E. models for input data with a small number of samples, and demonstrates a better sensitivity in D.E. analysis than its peers for the datasets with a few samples [8]. Finally, we demonstrate that the proposed feature selection method can also lead to meaningful network marker discovery for complex diseases.

## 2. Methods

Various feature selection algorithms are widely available for traditional omics data via different statistical tests [16]. However, most of these statistical tests based methods (e.g. *t-test*) can not apply to RNA-seq data directly, because they usually assume population data are normally distributed [17]. Some nonparametric statistical tests proposed for microarray data are available, but they are not widely employed in RNA-seq analysis probably because of their different generation mechanisms or differential analysis approaches [7, 12, 18].

On the other hand, traditional transform-based feature selection methods such as principal component analysis (PCA), nonnegative matrix factorization (NMF) or their variants can apply to RNA-seq data directly due to their purely data-driven characteristics, in which no distributions are assumed for input data [19, 20, 21, 22, 23, 24]. In fact, they transform input data into a subspace generated by principal components, or non-negative bases to seek meaningful linear combinations of features (genes). However, they face difficulties in ranking each gene because it is technically hard to distinguish an individual gene’s contribution to all genes’ linear combinations due to the nature of the linear or nonlinear transforms.

As such, it is believed that a desirable feature selection for RNA-seq data should satisfy the following criteria. First, it should be a data-driven method without prior data distribution assumption to prevent possible probabilistic modeling biases. Second, it should avoid evaluating each gene’s significance from the linear combinations of all genes in a subspace directly. Third, it should take consideration of the nonnegative characteristic of RNA-seq data instead of treating them as generic data to maintain locality in data analysis. Fourth, it should overcome the weakness of naive count-based filtering and contribute to following D.E. analysis by demonstrating robustness to depth and gene-length biases. That is, the feature selection algorithm should be more efficient than its other peers in identifying potential differentially expressed (DE) genes, rather than picking high-count genes or long genes only in feature selection.

We present a novel data-driven feature selection method: nonnegative singular value approximation (NSVA) that satisfies the criteria. It can be viewed as a special singular value decomposition (SVD) built upon Perron-Frobenius theorem, which is widely used in Google webpage ranking, to disclose novel findings for nonnegative data [25, 26, 27].

*Perron-Frobenius theorem.* Given matrix 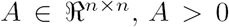, its largest eigenvalue 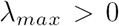 is always positive and its associative eigenvector *υ* is always positive, i.e. *υ* > 0. For any 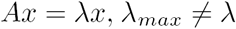, there exists at least one entry 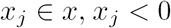.

### 2.1. Nonnegative singular value approximation (NSVA)

Given a nonnegative matrix *A* 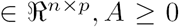 with a rank *r* = min(*n*,*p*), and its SVD decomposition 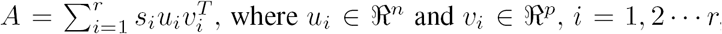, then, we have the following results,

1. Both *υ*_1_ and *υ*_1_ have only nonnegative entries, i.e., 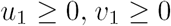.
2. The vectors *υ*_*j*_ and *υ*_*k*_ contain at least one negative entry when *j* ≥ 2, *j* = 1, 2 … *n*, and *k* ≥ 2, *k* = 1, 2 … *r*
3. Matrix A has the following first level singular value approximation:

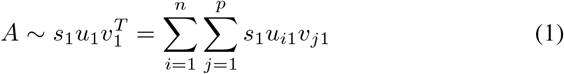

by dropping all *υ*_*i*_ and *υ*_*j*_, when *i,j* ≥ 2.

To prove nonnegative singular value approximation, we prove the following Perron-Frobenius extension theorem at first, which extends the results of the original theorem to nonnegative data.

#### 2.1.1. Lemma: Perron-Frobenius extension theorem

Given matrix 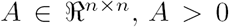, its largest eigenvalue *λ*_*max*_(*A*) ≥ 0 and its associative eigenvector *υ* is nonnegative, i.e. *υ*_*max*_ *≥* 0. *υ* > 0. For any For any 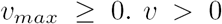, there exists at least one entry 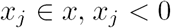.

Proof. We approximate *A* as a sequence of positive matrix *A*_*n*_ such as 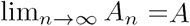. For example, if 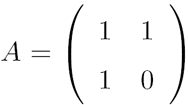, then it can be approximated as a sequence positive matrix 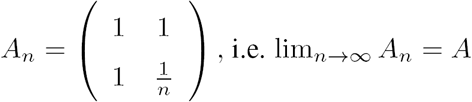.

It is clear that the characteristic equation of 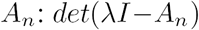 will also approximate the characteristic equation of 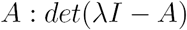 when *n →* ∞. As such,

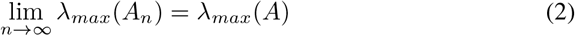

Since all *λ*_*max*_(*A*_*n*_) are positive, its limit should be nonnegative by the compactness of the sequence convergence, that is *λ*_*max*_(*A*) ≥ 0. Similarly, we normalize corresponding eigenvector *υ*_*n*_ of *λ*_*max*_(*A*_*n*_) such that 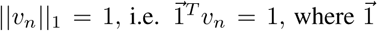 is a vector with all entries as 1. Thus, 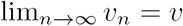.

Do limit for the following equation for the positive sequences,

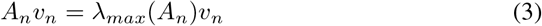

we have 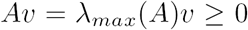. That is, *υ ≥* 0. Proceeding in the similar way, we can prove *υ >* 0. For any 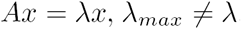, there exists at least one entry *x*_*j*_ ∈ *x, x*_*j*_ < 0.

#### 2.1.2. The proof of nonnegative singular value approximation

Suppose we have a nonnegative matrix 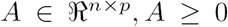, we assume *n ≥ p* without loss of generality. Then, 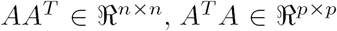, both are nonnegative semi-positive definite matrices.

By singular value decomposition (SVD): 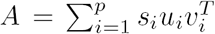, where *s*_*i*_ is the *i*^*th*^ singular value of *A*, it is easy to know *υ*_1_ and *υ*_1_ are the first eigenvectors of *AA*^*T*^ and *A*^*T*^ *A* respectively. That is, their corresponding eigenvalues are the first (largest) eigenvalue of *AA*^*T*^ and *A*^*T*^ *A*. The vectors *u*_*j*_, *j* = 2 … *n*, and *υ*_*k*_, *k* = 2 … *p* are the other eigenvectors of *AA*^*T*^ and *A*^*T*^*A* respectively.

Applying the Perron-Frobenius extension theorem, we have 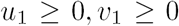. The other vectors *υ*_*j*_ and _*υ*_*k*, contain at least one negative entry. We only use the *υ*_1_ and *υ*_1_ to decompose *A* and drop *υ*_*j*_ and _*υ*__*k*_, for *j, k ≥* 2, we have 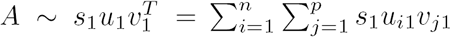.

#### 2.1.3. The biological meaning of nonnegative singular value approximation in RNA-seq analysis

It is noted that NSVA guarantees a purely additive decomposition of a nonnegative matrix along the first singular value direction 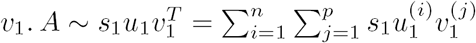. In fact, each nonnegative entry 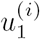 in *u*_1_ can be viewed as a corresponding coefficient of the row 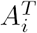, which represents the *i*^*th*^ gene of input RNA-seq data, in the one-dimensional “meta-sample space” spanned by all entries of 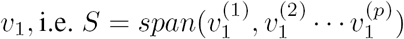 with a weight *s*_1_. Thus, from a single gene viewpoint, NSVA implies that each gene is approximated by the projection of its corresponding entry in vector *u*^1^ on the singular value direction 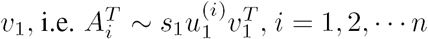.

Such an approximation makes it possible to rank each gene by using its coefficient in the meta-sample space, where each 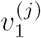 is the meta-sample corresponding to the original *j*^*th*^ sample, and 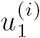 indicates the *i*^*th*^ gene 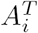’s contribution to all the meta-samples. It answers the following question: what’s a gene’s contribution to all meta-samples along with the first singular value direction? As such, it is natural to define 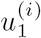 as a gene contribution score (GCS) to quantify its contribution to all meta-samples.

#### 2.1.4. Gene contribution scores (GCS)

*A gene contribution score (GCS)* measures a gene’s contribution to all samples of a RNA-seq dataset 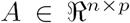 by evaluating its contribution to all meta-samples in a low dimensional space. The gene contribution score of the *i*^*th*^ gene to all samples is defined as 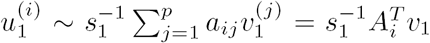 by applying nonnegative singular value approximation. It is clear that filtering genes according to their gene contribution scores is equivalent to filtering genes by their count variances by the nature of the GCS.

In fact, NSVA feature selection consists of two major steps. The first step conducts NSVA for input data and calculates GCS for each gene. The second step ranks all genes by their GCSs and selects the genes with large GCSs for the following D.E. analysis.

It is worthwhile to point out that the first singular value *s*_1_ is usually quite large for a RNA-seq dataset compared with the other singular values. we define a data variation explanation ratio as

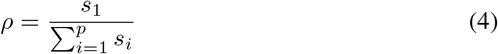

It is the ratio between the first singular value and the sum of singular values, to evaluate that the percentage of information can be represented in NSVA. The ratio actually represents the percentage of the data variances along the first singular vector direction. In fact, we have found that the ratio usually reaches at least *ρ* ≥ 60% or even 90% high for most RNA-seq datasets. For examples, the data variation explanation ratios of the three datasets in this study are 60:49%, 85.60% and 90.16% for the *Marioni, Prostate* and *Fly embryos* datasets respectively. In fact, the high data variation explanation ratios demonstrated by RNA-seq data guarantee the effectiveness of the first level singular value data approximation and the feasibility of the proposed feature selection algorithm.

### 2.2. NSVA-Seq: a data-driven differential expression analysis method

We propose a data-driven differential expression analysis method: NSVA-seq that employs NSVA to collect potential DE genes and compares each gene’s expression with those of remaining genes under a modified-fisher-exact-test (mFET) by computing exact p-values. Unlike other methods (e.g. DESeq), NSVA-seq avoids parameter estimation and tuning. Moreover, its average expression based hypothesis query under a contingency table can somewhat avoid the limitations of the existing D.E. analysis methods such that data with few number of replicates will not be ‘discriminated’ in D.E. analysis for its built-in disadvantage in parameter estimation or M-D odd ratio estimations[8, 11].

The NSVA-seq can be simply described as: given a normalized library, NSVA-seq applies the modified fisher exact test (mFET) to a set of genes selected by NSVA. Our modified fisher exact test (mFET) is described as follows.

The original fisher extract test is used to determine if there are non-random associations between two categorical variables [28]. In the modified fisher exact test, we query if a gene is differentially expressed by comparing it with all the remaining genes. Table 1 illustrates the mFET’s initial state: parameters a, b are the expression levels of a given gene *g* under the control and treatment conditions respectively. Similarly, parameters *c* and *d* are the expression levels of all remaining genes under the two conditions. All of these parameters are all non-negative integers initially.

**Table 1.**
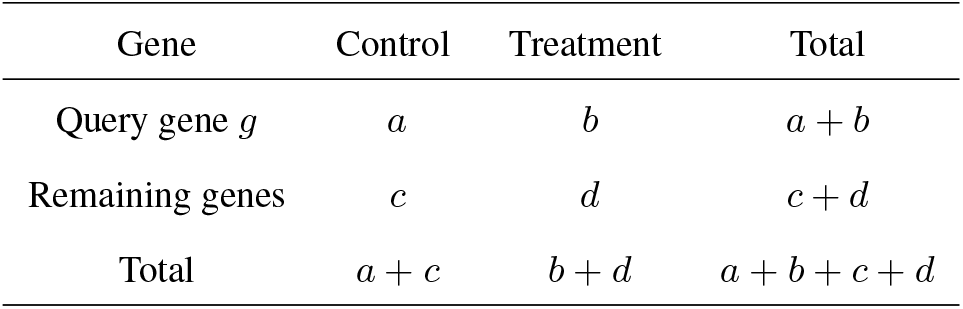
Modified fisher exact test (mFET)

| Gene | Control | Treatment | Total |
| --- | --- | --- | --- |
| Query gene $g$ | $a$ | $b$ | $a + b$ |
| Remaining genes | $c$ | $d$ | $c + d$ |
| Total | $a + c$ | $b + d$ | $a + b + c + d$ |

The null hypothesis **H**_0_ in context is to query whether a gene *g* has the same level expression as all the other remaining genes. The p-value of this hypothesis test can be calculated by a hypergeometric distribution as

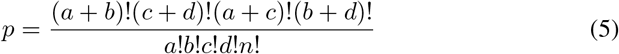

where *n* = *a* + *b* + *c* + *d*.

To avoid large computing complexities from the large or even huge values of gene count data, we apply a log transform to the equation and employ gamma function Γ(*n* + 1) to valuate *n*!. That is,

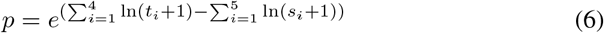

where 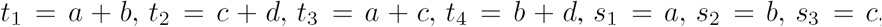 *s*_4_ = d, and *s*_5_ = *n*. In fact, we are able to extend the calculations to any nonnegative data instead of only nonnegative integers by the nature of the transformation. As such, the modified fisher exact test (mFET) can be reformulated as follows.

Given a RNA-seq data *X* 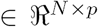, where each gene has two conditions control (C) and treatment (T), we have the following parameter specifications in the proposed mFET method:

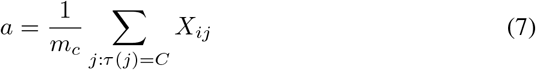

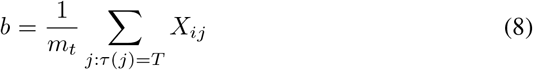

where *X*_*ij*_ is the expression level of the *i*^*th*^ gene of the sample *j*, *τ*(*j*) is the condition of the *j*^*th*^ sample, and *m*_*c*_ and *m*_*t*_ are number of samples in the control and treatment conditions respectively. Similarly, we have

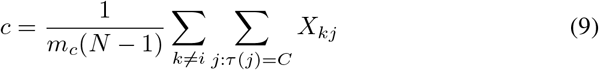

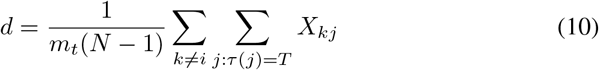

The proposed NSVA-seq provides more freedom in D.E. analysis than existing D.E. models. It can not only work well for normalized data, but also raw read count integer data. The proposed modified fisher exact test not only extends the input data domain of the original fisher exact test, but also generalizes its differential expression test for each gene by using the whole remaining gene expressions. As a nonparametric method, NSVA-seq does not need a parameter estimation process to find the mean and variance parameters of a specified distribution. As such, it somewhat provides a ‘fair’ D.E. analysis environment for those datasets consisting of few samples. It can be essentially important for clinical D.E. analysis, in which no enough samples are generally available [9, 13, 15]. On the other hand, it does not need to do M-D odd ratio comparisons as NOISeq for its more transparent D.E. analysis mechanism [11, 29].

## 3. Result

### 3.1. Datasets

We include three benchmark RNA-seq datasets in our experiments and their details are described as follows.

Marioni data originally consist of 32,000 genes across 14 samples after Illumina-supplied alignment algorithm ELAND. The samples are composed of two groups: the seven technical replicates from a kidney sample and another seven technical replicates from a liver sample, both of which are from a single human male [7]. This dataset is an important benchmark in normalization and D.E. analysis: it includes important gene length information for each gene compared to other RNA-seq datasets.

**Prostate data** consist of 17 million short reads and they were sequenced under the Illumina technology for two types of samples: four prostate cancer cells treated with androgen/DHT (DHT-treated), and three prostate cancer LNCap cells without DHT treatment (Mock-treated) [34]. We employed *Bowtie and SAMtools* to align its raw data with respect to the the human genome indexes (NCBI version 37), and obtained a nonnegative integer matrix with 4 DHT-treated and 3 Mock-treated samples across 23,068 genes [5, 30].

**Fly embryos data** consist of 17,605 genes across four samples. The four samples are composed of two biological replicates at conditions “A” (treated) and “B” (control) respectively. This dataset consists of only four samples and usually demonstrate some disadvantage in the existing D.E. models that require relatively more samples to complete parameter estimation [8, 13, 15].

### 3.2. DESeq analysis with nonnegative singular value approximation (NSVA) feature selection

To verify the effectiveness of the proposed feature selection, we firstly combine it with DESeq model, which is a typical parametric D.E. analysis model, to answer the query: ‘what will happen to DESeq analysis when NSVA feature selection is applied to input data?’ Figure 1 evaluates the performance of three NSVA-selected gene-sets consisting of 2,000, 3,000, and 5,000 genes and original data under DESeq. The false discovery ratio (FDR) cutoff was uniformly chosen as 0.001 in our experiments. Each horizontal and vertical axis in the subplots represent the mean expression of each gene and its corresponding log_2_ fold changes under two different conditions respectively. The differentially expressed (DE) and non-DE genes are represented by red and blue markers respectively.

**Figure 1:**
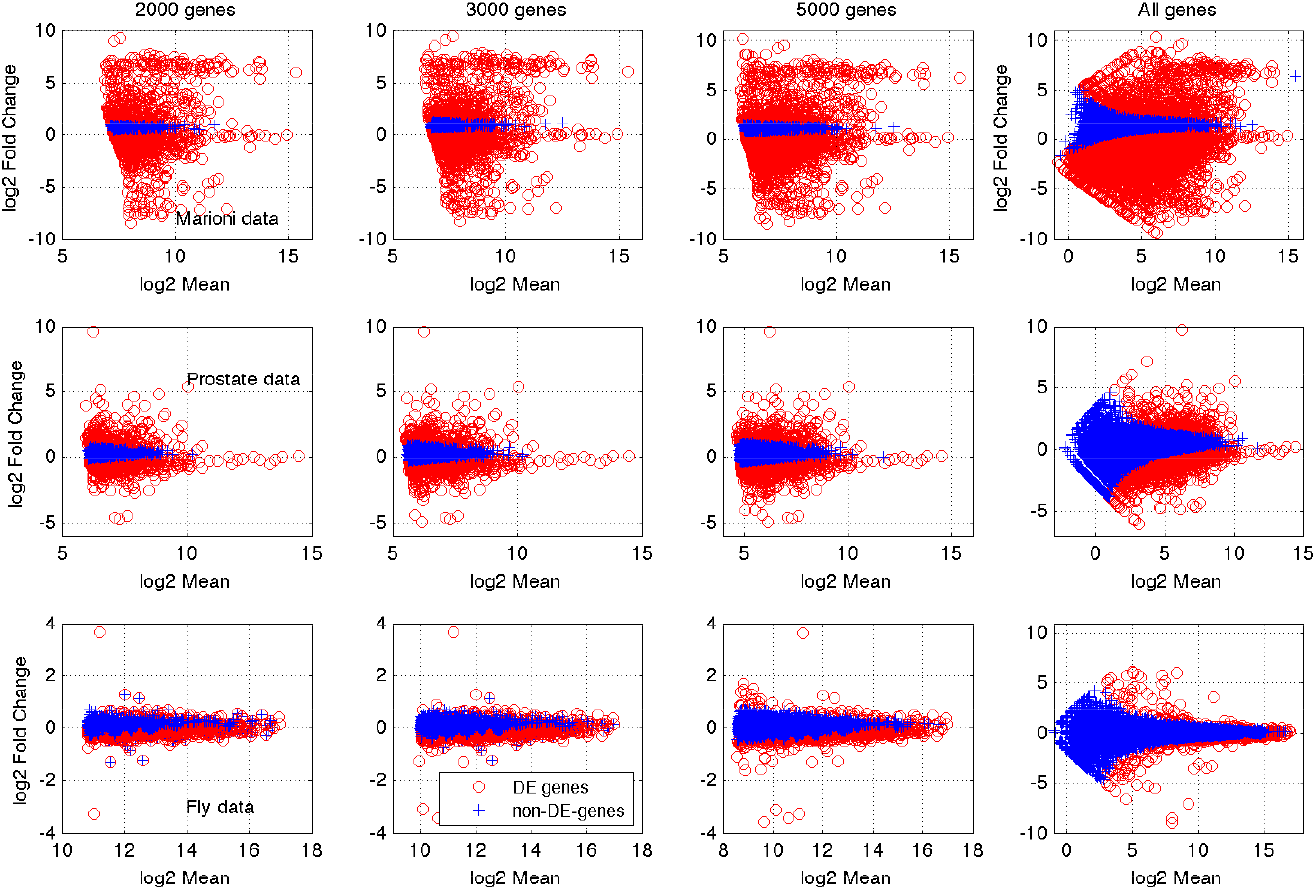
The scatter plots of normalized data mean versus *log*_2_ fold change for original data and different NSVA-selected gene sets under DESeq on Marioni and Prostate datasets. The D.E. and non-D.E. genes are represented by red and gray markers respectively. The non-D.E. genes dropped remarkebaly in DESeq analysis when NSVA is applied to filter more genes.

Interestingly, The non-D.E. genes seem to drop remarkably under DESeq when NSVA is applied to each dataset. It indicates that the proposed NVSA feature selection demonstrates a good sensitivity to filter those non-differentially expressed (non-DE) genes for each dataset by picking the genes with large gene contribution scores (GCS). In other words, NSVA seems to be able to select more potential DE genes, which have more contributions to variance on behalf of the first singular value direction. Such a feature selection makes the following DESeq analysis more focused on the potential DE genes and contributes to decreasing false positives in D.E. analysis. Such a result suggests that the proposed feature selection can enhance D.E. analysis by picking meaningful genes.

#### 3.2.1. The impacts of NSVA feature selection on the DESeq model

In addition to comparing NSVA-DESeq with DESeq from a performance stand point, we have the following findings about NSVA’s impacts on the DESeq model itself.

First, we have found that NSVA feature selection will contribute to a better size factor estimation in normalization because of filtering outliers (e.g., genes with very low counts). The size factor *s*_*j*_ of a sample *j* in DESeq model is a normalization factor to make the sample, which may be subject to different sequencing depth, comparable with the others. However, the size factors actually relies on a pseudo-reference sample, which is a virtual sample consisting of geometric means of all genes [8]. Filtering the outliers will prevent their geometric means from being entries of the pseudo-reference sample, which will cause the size factor estimation to be closer to the ‘truth’ and mitigate the bias caused by the sequencing depth.

Second, NSVA feature selection makes the parameter estimations of *u*_*ij*_ and 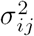, which are mean and variance parameters of gene *x*_*ij*_ (the *i*^*th*^ gene on the *j*^*th*^ sample), under a negative binomial (NB) distribution, more robust. This was partially because both parameter estimations were strongly dependent on the estimation of the size factor *s*_*j*_ [8]. More interestingly, we found that the smooth function used by the DESeq method to model the dependence of the raw variance on the mean was fitted much better using NSVA-selected genes than the all genes in the local regression.

Figure 2 illustrated the means and raw variances of seven liver samples in the *Marioni* data, and the fitting of the raw variances with respect to the means (red lines) using all 15,514 genes and only 2000 NSVA-selected genes, in the NW and NE plots respectively. The similar results can be also found for the *Fly embryos* data with 4 samples but more than 17000 genes, in the SW and SE plots of Figure 2. It is clear that the fits under the NSVA-selected genes are much better than those under all genes. More importantly, such a good fitting contributes to more accurate variance parameter estimation in DESeq, which will enhance the accuracy of the following hypothesis test in the differential expression call.

**Figure 2:**
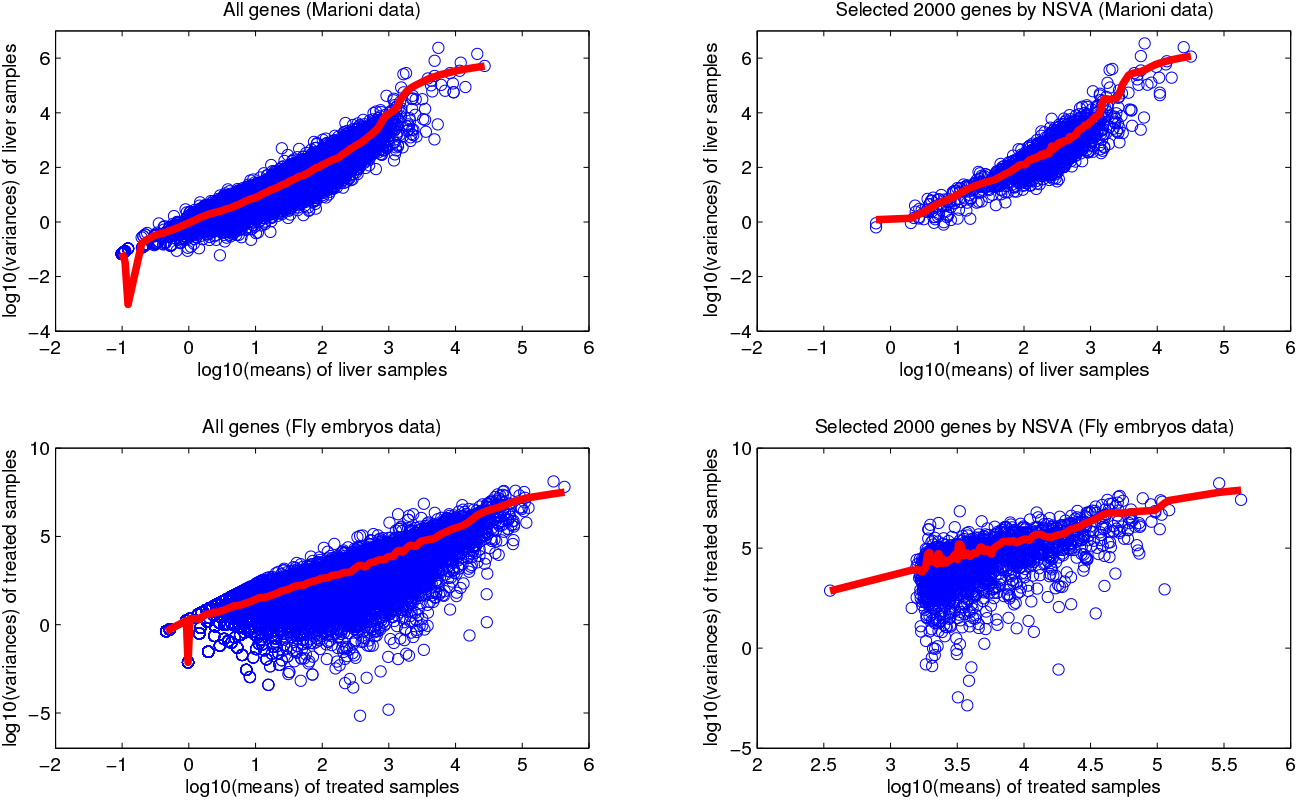
The plots of the sample variances with respect to the means of 7 *Liver* samples of the *Marioni* dataset under all 15,514 genes and NSVA-selected 2000 genes and the similar plots for the *Fly embryos* data. The red lines represent the fits of raw variances with respect to the means of the *Liver (Fly embryos)* samples respectively.

Last, NSVA feature selection contributed to decreasing the complexity of the hypothesis test in DESeq due to the fact that a lot of genes were filtered by NSVA, which actually avoids quite a lot computing burden because calculating the p-value for each gene in DESeq requires to enumerate all possible count sum combinations of two conditions (e.g., treated vs untreated) from a given total count sum [8].

### 3.3. Compare nonnegative singular value approximation with peer methods

We further compare our NSVA feature selection with other peer methods to demonstrate its advantage in picking potential DE genes. These methods include count-based naive feature selection (NFS), principal component analysis (PCA), nonnegative matrix factorization (NMF), signal-noise-ratio (SNR), and geometric signal-noise-ratio (GSNR) [20, 19, 31, 32, 33]. All the five comparison methods are data-driven methods as NSVA.

In fact, the count-based naive feature selection (NFS) is just the widely used low-count filtering method. PCA and NMF both belong to variance-based feature selection methods as NSVA though they use different variance metrics in feature selection. SNR and GSNR belong to signal-noise based feature selection method that rank each gene via comparing signal-noise ratios.

Count-based naive feature selection (NFS). As a coverage-based feature selection method, NFS filters the genes with low counts and keeps those with high counts before D.E. analysis. It selects all genes ≥ the median gene count of the input data, sorts all genes according to its coverage, i.e., the sum of its counts, and picks the top-ranked genes before D.E. analysis [8, 13].

#### 3.3.1. Principal component analysis (PCA)

As a variance-based feature selection method, PCA ranks each gene by using the 2-norm of its projection in the subspace spanned by the first three principal components (PCs) by measuring the gene’s contribution to the three most significant PCs [19, 32]. It is noted that the three major PCs usually have a quite high variance explained ratio (e.g., 99%) for each dataset. The PCA feature selection consists of the following three steps.

The first step conducts PCA for input data *X* 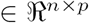 to obtain the principal component (PC) matrix: *U ← princomp*(*X*), and projected data to the first three PCs, i.e. 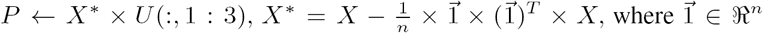 is a vector with all entries ‘1’. The second step calculates the 2-norm for the projection data of each gene in the subspace spanned by the three 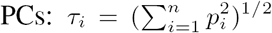, where *p*_*i*_ is the *i*^*th*^ row of the projection matrix 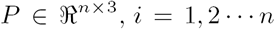. Finally, the third step sorts the genes according to *τ*_*i*_ and selects the top-ranked genes.

#### 3.3.2. Nonnegative matrix factorization (NMF)

Similar to PCA, NMF is a variance-based feature selection but requires the non-negativity of input data [20]. Given an input RNA-seq data 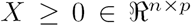, NMF conducted the following decomposition: *X* ~ *WH* at rank *p –* 1 firstly, where 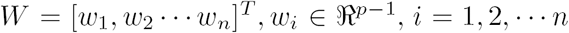, and 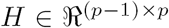, Then, 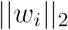 is used to rank the *i*^*th*^ gene’s contribution to the whole data variance. Finally, the top-ranked genes were selected by sorting the values of 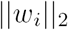.

#### 3.3.3. Signal-Noise ratio (SNR) and geometric signal-noise ratio (GSNR)

This data-derive feature selection method ranks each gene by the ratio of gene mean and standard deviation: 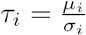, where *µ*_*i*_ and *σ*_i_ are estimated as 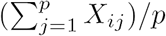 and 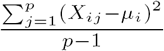 respectively for given *X* 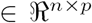. These genes with large SNR values are believed to be more meaningful genes. It is noted that the infinite SNR values are set as zeros automatically in our feature selection [31].

Different to the SNR feature selection, this method ranks each gene by using the geometric signal-noise ratio to rank each gene. GSNR is defined as the ratio of the geometric mean and geometric standard deviation as 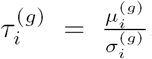, where 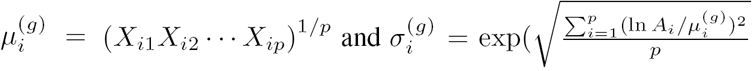 respectively for given *X* 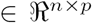 These genes with large 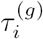 values are believed to be more meaningful genes in feature selection [33].

#### 3.3.4. NSVA is robust to depth and gene-length biases

We need to answer the following two questions: 1) Is NSVA more efficient than its peers in identifying potential DE genes? 2) Is NSVA a depth-dependent feature selection method, where high-count genes are more likely to be identified as DE genes?

To answer the queries, we compare proposed NSVA with its peers on two measures under DESeq analysis: *DE ratios* and *DE gene median counts*. The DE ratio refers to the ratio between the number of DE genes identified by a differential expression analysis *Ω*, which is employed as DESeq analysis in this context, and the total number of genes: *N* of the input data, where ε is a significant level cutoff (e.g. 0.01) and *θ* is a feature selection method employed before differential expression analysis, namely,

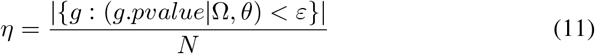

It measures the efficiency of a feature selection method. An efficient feature selection method *θ* should produce a high DE ratio for a dataset under a specified DE analysis *Ω*. The DE gene median count τ is the read count median among all DE genes

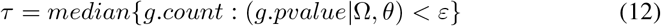

under Ω and *θ*. A feature selection method would be a depth-dependent method, provided it had high median counts for DE genes.

Figure 3 demonstrates the DE ratios and DE gene median counts of proposed NSVA and its five peers on the *Marioni* and *Prostate* data, when 2000, 3000, 5000, and 8000 genes are selected in feature selection [7, 34]. Interestingly, the results suggest that NSVA is a more efficient method compared with its peers. It achieves the highest DE ratios for each case among all feature selection methods. The NFS feature selection performs a little bit better than NMF, PCA and GSNR. It indicates that complicate transform-based feature selection methods (e.g. NMF) may not contribute to DE analysis. SNR has the worst DE ratios. It indicates that simple feature selection methods like SNR can not contribute to DE analysis either.

**Figure 3:**
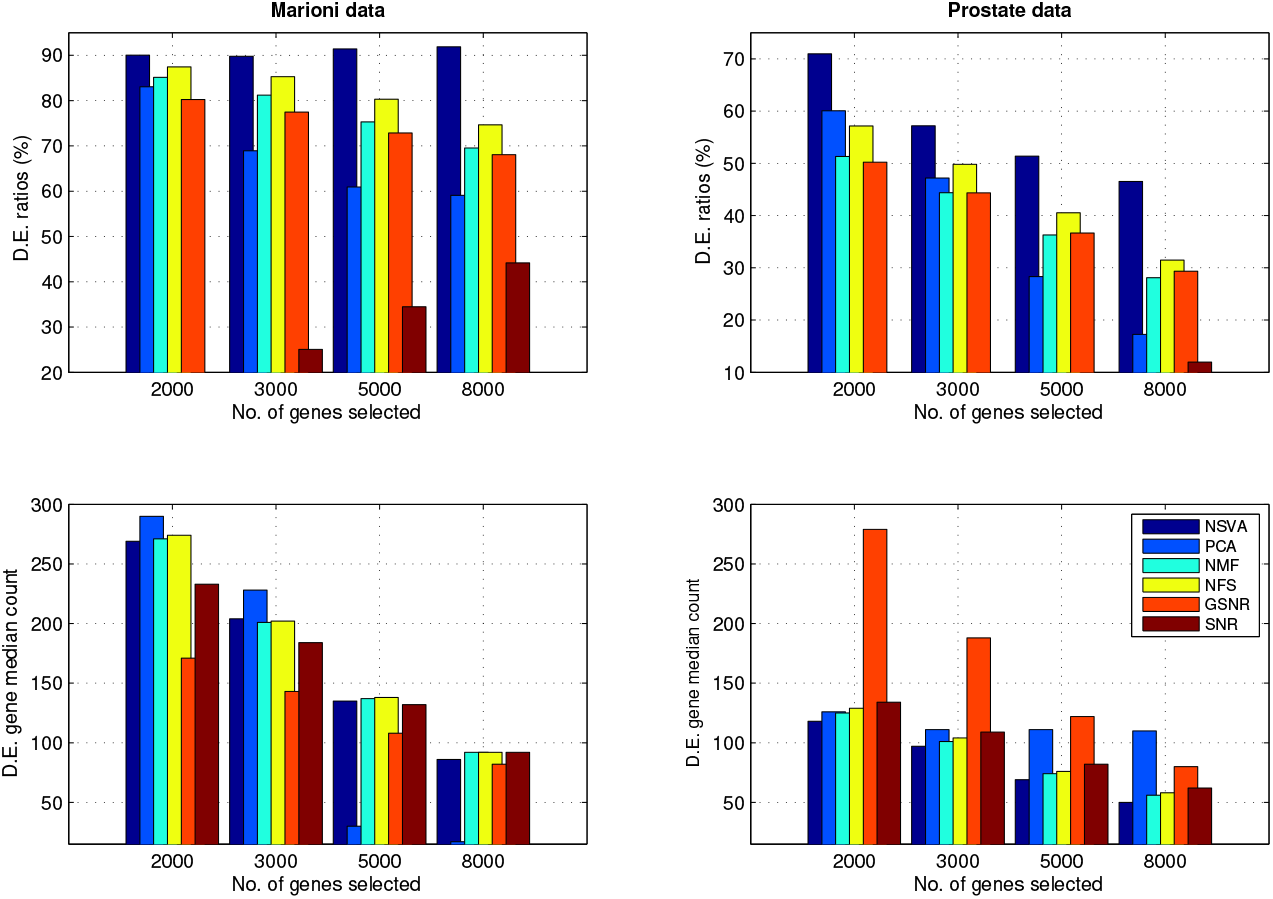
The comparisons of DE ratios and DE gene median counts for NSVA and its peers under DESeq analysis on two datasets. The proposed NSVA feature selection demonstrates strong advantages in selecting potential DE genes than its competing methods by producing highest DE ratios. The DE gene median counts of NSVA are generally lower than those of other peers for the two datasets.

In addition, NSVA has the shortest median DE gene count values than all the other methods for *Prostate* data. On the other hand, the DE gene median counts of NSVA are generally lower than those of PCA and NFS, equivalent to that of NMF, and higher than those of GSNR and SNR for the *Marioni* data. Those results strongly suggest that NSVA should not be a depth-dependent feature selection method like NFS.

##### Does NSVA only pick long genes in feature selection?

That is, NSVA can contribute to avoiding gene-length bias in RNA-seq analysis? To answer this query, Figure 4 compares the gene length medians of the genes selected by the six feature selection methods and the DE genes among the selected ones for the *Marioni* data. The other two datasets have no gene length information available.

**Figure 4:**
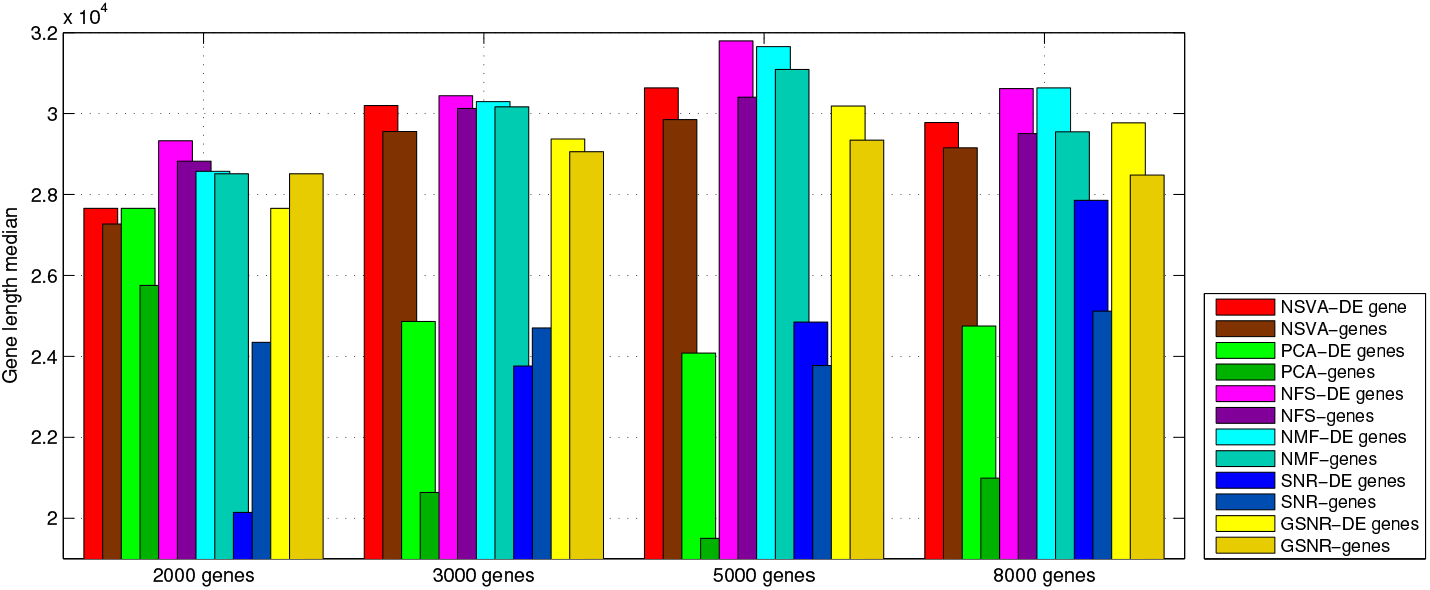
The comparisons of the gene length medians of the genes selected by different feature selection methods and DE genes among the selected genes for the Marioni data. The DE genes generally have longer gene length than those selected genes from almost all methods. The NSVA-selected genes and their DE genes are shorter than those from their peers like NFS, and NMF.

It is interesting to see that the DE genes generally have longer gene length than those selected genes from almost all methods except SNR. Such a result is consistent with the consensus that long genes are more likely to be selected as DE genes in RNA-seq D.E. analysis. Furthermore, NSVA-selected genes and its DE genes are shorter than those from NFS and NMF, but longer than those from PCA and SNR. For example, The median gene lengths of NSVA-selected genes in all the four gene-selection cases are higher than the DE gene median length (26,445 bp) of all genes for the *Marioni* data [7]. Furthermore, its DE gene median length has reached 27,659 bp on the 2,000 gene selection case, which is much lower than that of NFS (29,328 bp) and NMF (28, 513 bp). GSNR has an almost same level gene median length as NSVA, but it has relatively lower DE ratios than NSVA.

Table 2 compares the six methods on behalf of DE ratios, DE gene count median, gene-median-length and DE gene-median-length. The gene-median-length and DE gene-median-length refer to the gene median length of NSVA-selected genes and DE genes among the NSVA-selected genes respectively. It is clear that NSVA is more efficient than its peers in identifying potential DE genes for its highest DE ratios. NSVA also demonstrates it is not a depth-dependent feature selection method, where high-count genes are more likely to be identified as DE genes for its low DE gene-count-medians. Furthermore, NSVA selects ‘median-long’ genes instead of only long genes or short genes from the gene median length of the NSVA-selected genes and DE genes among them.

**Table 2.**
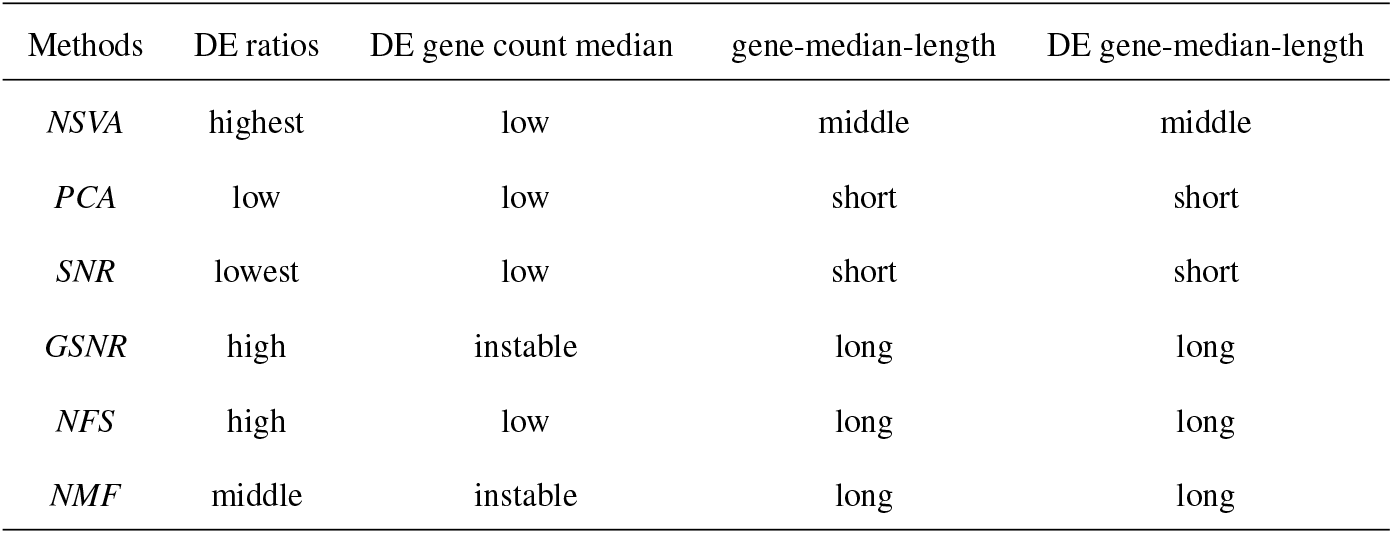
The comparisons of six feature selection methods.

| Methods | DE ratios | DE gene count median | gene-median-length | DE gene-median-length |
| --- | --- | --- | --- | --- |
| *NSVA* | highest | low | middle | middle |
| *PCA* | low | low | short | short |
| *SNR* | lowest | low | short | short |
| *GSNR* | high | instable | long | long |
| *NFS* | high | low | long | long |
| *NMF* | middle | instable | long | long |

As such, NSVA seems to be the best one among the six feature selection methods for its robustness in depth bias and gene length biases. It not only avoids only picking those long genes or high-count genes like NFS/NMF, but also the short genes or low-count genes as PCA/SNR by considering its high DE ratios and low DE gene medians counts. That is, it can contribute to picking potentially DE genes and decreasing the false positives in DE analysis.

### 3.4. Nonnegative singular value approximation for non-parametric D.E. models

We further apply NSVA to non-parametric D.E. method NOISeq to demonstrate its effectiveness in differential expression analysis. The NOISeq employs two statistics: *M* and *D* to compare these to the noise distribution to determine whether the expression is statistically significant. The *M* and *D* values measure the *log*_2_ fold change and *log*_2_ absolute expression difference between conditions. From this comparison, NOISeq produces the probability value of their odd ratio that, when compared to a threshold number (*q*), which is set as 0.8 in our experiment, determines whether the gene is actually differentially expressed [11].

#### Applying NSVA to NOISeq

Like DESeq, NOISeq demonstrates the increase of DE ratios in D.E. analysis when using NSVA feature selection [8, 11]. Figure 5 illustrates the scatterplots of plotted (M,D) values (M-D plots) produced from our NOISeq method for all the datasets. The M-D plot is essentially to impose an M-plot on a D-plot, which is similar to volcano plot in traditional microarray analysis [35].

**Figure 5:**
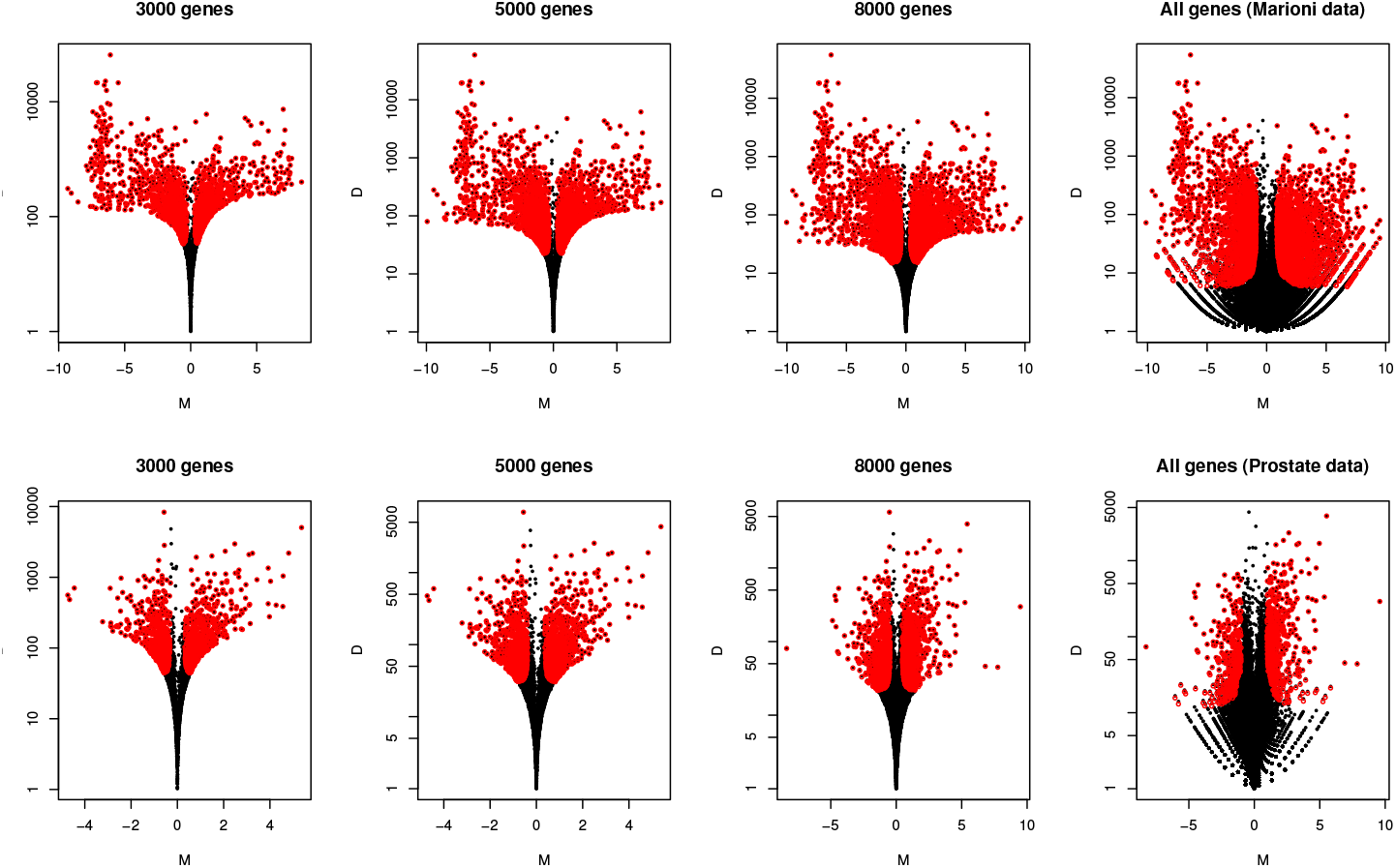
The comparisons of the M-D plots of NSVA-selected genes and the original *Marioni* and *Prostate* data in *NOISeq* analysis, where DE and non-DE genes are indicated by red and black dots respectively. The non-DE genes drop remarkably in NOISeq when NSVA is applied to filter more genes

The plots compare the M-D plots of the top 3000, 5000, and 8000 NSVA-selected genes, along with the original dataset under NOISeq respectively. The red and black dots represent differentially expressed genes, and non-differentially expressed genes. It is clear that non-DE genes drop remarkably when more genes are filtered by NSVA for the two datasets. In other words, the corresponding DE ratio would increase for each selected gene set under such a feature selection. Obviously, such a result is consistent to the previous results from applying NSVA to the parametric model: DESeq [8]. It further indicates that such a feature selection can enhance D.E. analysis by picking meaningful genes for both parametric and non-parametric D.E. analysis models.

Table 3 compares the DE ratios of NSVA-selected datasets and original data under NOISeq. It is clear to see that DE ratios increase for all three datasets when more genes are filtered in NSVA-feature selection. For example, when the 2000 most significant genes are selected from the *Marioni* dataset, 81.9% of those genes are determined to be differentially expressed. But the DE ratios of the original dataset without feature selection has only 17.52%. On the other hand, the DE ratio of the original *Fly* dataset is only 0.32%, but such a ratio reaches 2.4% when the 2000 most significant genes are selected in feature selection.

**Table 3.** DE ratios of NSVA-selected datasets and original data under NOISeq.

| Selected genes | DE ratios (*Fly* data) | DE ratios (*Prostate* data) | DE ratios (*Marioni* data) |
| --- | --- | --- | --- |
| 2000 genes | 2.4% | 54.45% | 81.90% |
| 3000 genes | 1.87% | 47.27% | 78.27% |
| 5000 genes | 1.04% | 37.40% | 72.56% |
| Total data | 0.32% | 22.92% | 17.52% |

We also conduct naive feature selection (NFS) for NOISeq by removing all genes with count < 10. However, it can’t achieve good DE rations as NSVA. For example, the DE ratio is only 42.92% for 5606 NFS-selected genes for the *Marioni* dataset, but the DE ratio under the 5000 NSVA-selected genes is 72.56%. In addition, the DE ratio is only 30.93% for 1809 NFS-selected genes for the *Prostate* dataset, but the DE ratio under the 2000 NSVA-selected genes is 54.45%. Such results again indicate the proposed feature selection performs better than naive feature selection (NFS) in selecting meaningful genes.

### 3.5. Compare NSVA-seq with peer D.E. analysis models

To demonstrate the effectiveness of proposed NSVA-seq, we apply it to the gene set consisting of top 10% genes ranked by NSVA from each dataset normalized by DESeq normalization [8][10]. Then, we compare its adjusted p-value distributions with those of four peer methods: NSVA-DESeq, NSVA-edgeR and NSVA-NOISeq and mFET, where mFET is applied to the original normalized data. It should be noted that the notations NSVA-DESeq/edgeR/NOISeq refers to applying DESeq/edgeR/NOISeq analysis to the NSVA-selected genes respectively. We employ Benjamini-Hochbert (BH) procedure to adjust all p-values under a FDR 0.01 in such a comparison.

Figure 6 illustrates the scale of the adjusted p-values from NSVA-seq is in a quite small range compared with those of the others. Such a result strongly suggests that NSVA-seq is more sensitive to identify those genes with quite small adjusted p-values in differential expression analysis than the others. For example, almost all adjusted p-values are less than 0.03 that indicates these identified DE genes have a strong p-value support. In contrast, mFET generates a large amount of non-DE genes without NSVA feature selection for each dataset. It implies that NSVA tends to pick DE genes in feature selection, which directly contributes to the high DE ratios of NSVA-seq. As such, NSVA-seq has a much lower false positive ratio than mFET.

**Figure 6:**
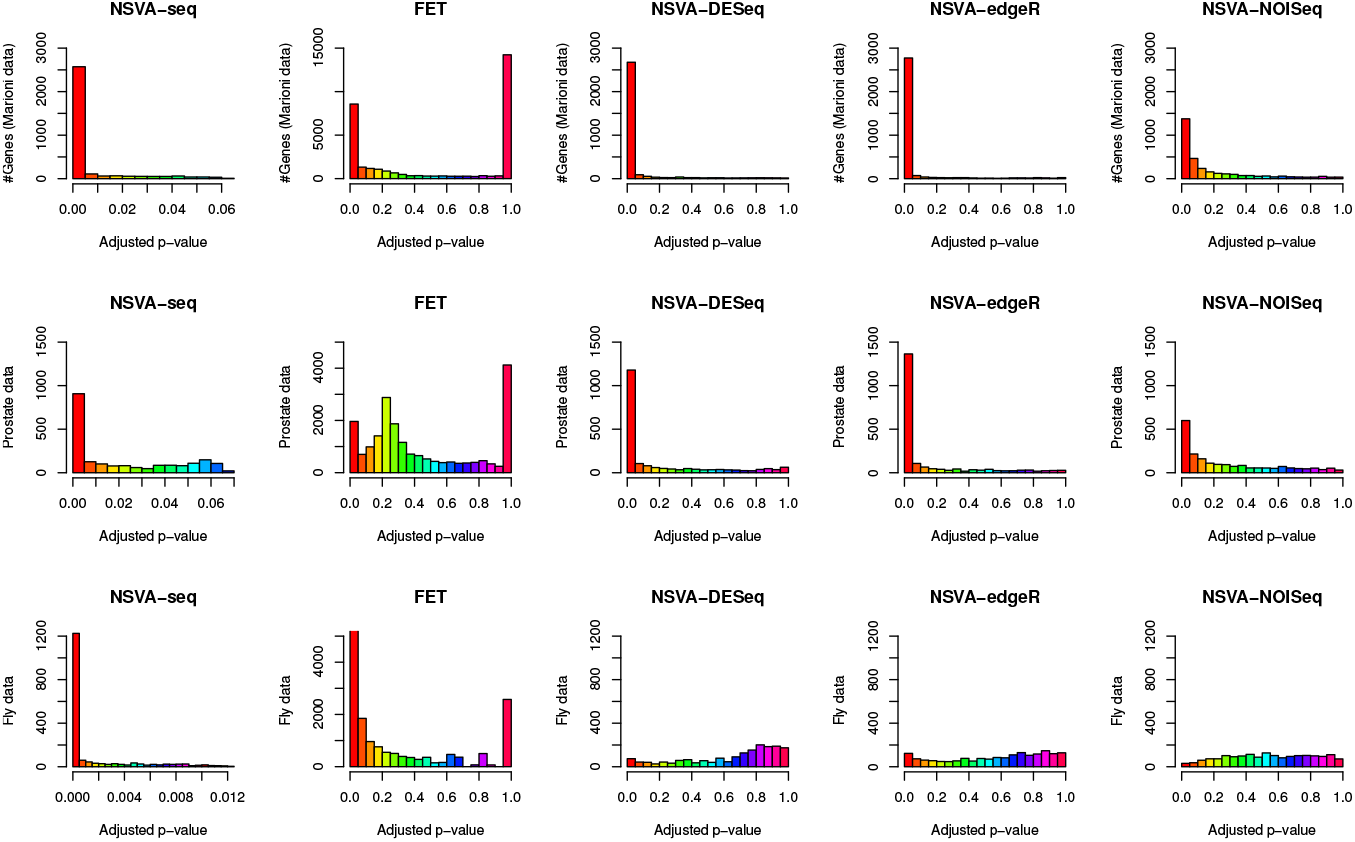
The comparisons of the adjusted p-value distributions of NSVA-Seq, mFET, NSVA-DESeq, NSVA-edgeR, and NSVA-NOISeq. NSVA-seq demonstrates a conservative D.E. analysis for the *Marioni* and *Prostate* datasets. But it overcomes the weakness of its peers in D.E. analysis of the *Fly embryos* dataset with few samples.

On the other hand, NSVA-DESeq and NSVA-edgeR have similar distribution patterns due to the same underlying assumption on the count data distribution of the DESeq and edgeR models. It is even hard to claim the advantage of DESeq than edgeR in D.E. analysis under NSVA-feature selection [8, 13, 15]. Such a result further implies that power of NSVA in selecting potential DE genes.

Interestingly, NSVA-seq seems to be more conservative in D.E. analysis than NSVA-DESeq and NSVA-edgeR for the *Prostate* and *Marioni* datasets. However, it actually identifies more DE genes for the *Fly embryos* dataset that has only 4 samples than the other methods, which seem to identify almost all genes as non-DE genes. This is because the datasets with few samples have some disadvantage in estimating accurate mean and variance parameters parameters for parametric D.E. analysis models like DESeq and edgeR [8, 13]. For example, a dataset with few samples may cause difficulties for the local fit procedure in the DESeq model [8, 15]. On the other hand, too small sample size can lead to low odd-ratios of *M* and *D* in NOISeq due to the lack of replicates and the likelihood to miscount meaningful expression signals as noise [11]. However, our NSVA-seq avoids parameter estimation or *M-D* odd ratio comparisons by comparing each gene’s average expression with a set of selected genes’ average expressions. It mitigates the side-effect from the small number of samples and provides a fair D.E. analysis environment for RNA-seq datasets. It is worthwhile to point out that similar results can be obtained when TMM is employed as the procedure in data normalization rather than DESeq normalization [8, 9].

Furthermore, NSVA-seq has a more transparent D.E. analysis mechanism than NOISeq though both are nonparametric D.E. analysis models. Its modified FET based D.E. analysis under NSVA-feature selection is more direct than NOIseq that relies on M-D odd ratio comparisons [9]. However, NSVA-seq demonstrates advantages in overcoming the weakness of the existing D.E. models in handling datasets with few samples, besides more conservative D.E. analysis results for other datasets. Such a characteristic can be essentially useful for clinical D.E. analysis, in which no enough samples are generally available [9, 13, 15].

#### 3.5.1. Nonnegative singular value approximation based biomarker discovery, a case study

We further demonstrate the effectiveness of NSVA in biomarker discovery from by using jActiveModule to search active subnetwork modules [37]. We use the top-ranked 2000 genes with smallest probability values under NSVA-DESeq to find possible biomarkers for the *Prostate* data, in which input dataset consisting of 5000 genes selected by NSVA from the original *Prostate* data. We have found that there are several networks with varying active path scores of: 4.97, 5.22, 5.24, 5.29, and 5.98. We use the module with the highest score: 5.98, as our network marker that has 179 nodes and 630 edges.

Figure 7 illustrates the network marker by high-lightening those genes with most protein-protein interactions. Although detailed analysis of such a network marker is beyond the scope this study, we would like to analyze the genes with the largest interactions in the network marker. YWHAE, TARDBP, and CALM1 are the three genes with the most interactions among the network marker. It is interesting to see that all of them have much closer relationships with prostate cancer. For example, YWHAE belongs to the 14-3-3 family of proteins which mediate signal transduction by binding to phosphoserine-containing proteins and has been reported to have express in prostate cancer [38]. Furthermore, it was reported to interact with YWHAZ, a widely known biomarker of prostate cancer [39, 40]. In addition, TARDBP has been found to have multiple functions in transcriptional repression, pre-mRNA splicing and translational regulation. It was reported as one of biomarkers to distinguish prostate cancer from benign prostatic hyperplasia in patients[41]. Moreover, CALM1’s mutation was reported to connect with prostate cancer and was one of verifiable biomarkers of prostate analysis using urinary shotgun proteomics [42, 43]. Such a meaningful biomarker capturing indicates the usefulness of our network marker though more detailed analysis can be done for this network marker to retrieve more comprehensive information. Alternatively, it demonstrate the effectiveness of NSVA in D.E. analysis and biomarker discovery.

**Figure 7:**
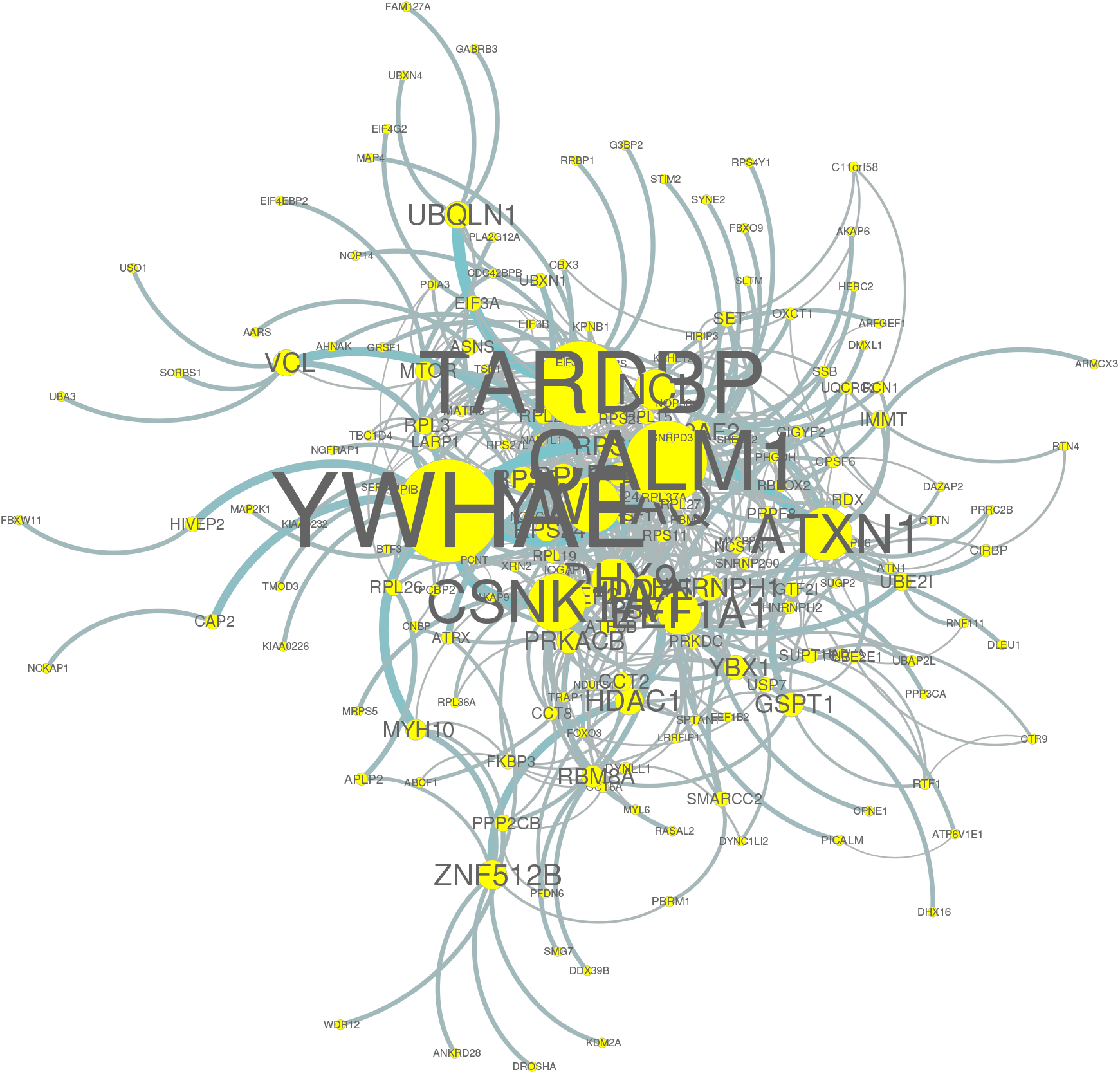
The network marker of the *Prostate* dataset based on the 2000 most significant genes from NSVA-DESeq. The top three gene with most interaction are YWHAE, TARDBP, an CALM1.

## Conclusion

In this study, we propose a novel NSVA feature selection and NSVA-seq differential expression analysis method for RNA-seq data. The NSVA feature selection is rooted in a rigorous mathematical result from singular value decomposition for non-negative RNA-seq read count data. The proposed NSVA-based feature selection algorithm demonstrates robustness to depth and gene length robustness by overcoming the weakness of widely used naive count feature selection (NFS). It demonstrates advantages in picking meaningful potential DE genes for different D.E. analysis models by enhancing the efficiency of D.E. analysis by comparing with its five peer feature selection methods.

As a data-driven D.E. analysis, NSVA-seq provides more freedom in D.E. analysis by allowing both original count data and normalization data in D.E. analysis. It not only avoids the parameter estimation process, but also provides a more direct and transparent nonparametric D.E. analysis, which contributes to easy understanding and implementation. More importantly, it overcomes the limitations of the existing D.E. analysis models by providing a fair D.E. analysis for those datasets with few samples besides producing a relatively conservative D.E. analysis for the other datasets. Furthermore, the biomarker discovery results demonstrate the effectiveness of NSVA in capturing meaningful genes, and its positive impacts on D.E. analysis and meaningful gene marker capturing for complex diseases.

However, how to achieve an optimal feature selection for the sake of robust differential expression analysis remains a challenge for this method. We are employing information measures such as mutual information or entropy to explore its potential in NSVA feature selection [44]. Moreover, we are interested in conducting novel pathway analysis for the network marker obtained in this study to dig more knowledge and further enhance its repeatability and validity [45].

In addition, we are applying NSVA and NSVA-seq to RNA-seq datasets retrieved from TCGA portal, which are a type of structured big data, to further investigate the effectiveness of our methods [46, 47]. Those datasets can be high-dimensional imbal-anced data (HDI): high-dimensional data with skewed label distributions. They usually bring hard time in disease diagnosis when there is no feature selection done [48]. We are interested in investigating the impacts of NSVA feature selection on such data to further explore its potential in disease diagnosis.

## Acknowledgement

Author sincerely thanks Dr. Wentian Li for his invaluable suggestions and comments for this work.

